# Epigenome-wide meta-analysis of blood DNA methylation and its association with subcortical volumes: findings from the ENIGMA Epigenetics Working Group

**DOI:** 10.1101/460444

**Authors:** Tianye Jia, Congying Chu, Yun Liu, Jenny van Dongen, Nicola J Armstrong, Mark E. Bastin, Tania Carrillo-Roa, Anouk den Braber, Mathew Harris, Rick Jansen, Jingyu Liu, Michelle Luciano, Anil P.S. Ori, Roberto Roiz Santiañez, Barbara Ruggeri, Daniil Sarkisyan, Jean Shin, Kim Sungeun, Diana Tordesillas Gutiérrez, Dennis van’t Ent, David Ames, Eric Artiges, Georgy Bakalkin, Tobias Banaschewski, Arun L.W. Bokde, Henry Brodaty, Uli Bromberg, Rachel Brouwer, Christian Büchel, Erin Burke Quinlan, Wiepke Cahn, Greig I. de Zubicaray, Tomas J. Ekström, Herta Flor, Juliane H. Fröhner, Vincent Frouin, Hugh Garavan, Penny Gowland, Andreas Heinz, Bernd Ittermann, Neda Jahanshad, Jiyang Jiang, John B. Kwok, Nicholas G. Martin, Jean-Luc Martinot, Karen A. Mather, Katie L. McMahon, Allan F. McRae, Frauke Nees, Dimitri Papadopoulos Orfanos, Tomáš Paus, Luise Poustka, Philipp G. Sämann, Peter R. Schofield, Michael N. Smolka, Lachlan T. Strike, Jalmar Teeuw, Anbupalam Thalamuthu, Julian Trollor, Henrik Walter, Joanna M. Wardlaw, Wei Wen, Robert Whelan, Liana G. Apostolova, Elisabeth B. Binder, Dorret I. Boomsma, Vince Calhoun, Benedicto Crespo-Facorro, Ian J. Deary, Hilleke Hulshoff Pol, Roel A. Ophoff, Zdenka Pausova, Perminder S. Sachdev, Andrew Saykin, Margaret J. Wright, Paul M. Thompson, Gunter Schumann, Sylvane Desrivières

**Author notes:** Corresponding author: Sylvane Desrivières. These authors contributed equally: Gunter Schumann and Paul M. Thompson.

## Abstract

DNA methylation, which is modulated by both genetic factors and environmental exposures, may offer a unique opportunity to discover novel biomarkers of disease-related brain phenotypes, even when measured in other tissues than brain, such as blood. A few studies of small sample sizes have revealed associations between blood DNA methylation and neuropsychopathology, however, large-scale epigenome-wide association studies (EWAS) are needed to investigate the utility of DNA methylation profiling as a peripheral marker for the brain. Here, in an analysis of eleven international cohorts, totalling 3,337 individuals, we report epigenome-wide meta-analyses of blood DNA methylation with volumes of the hippocampus, thalamus and nucleus accumbens (NAcc) –three subcortical regions selected for their associations with disease and heritability and volumetric variability. Analyses of individual CpGs revealed genome-wide significant associations with hippocampal volume at two loci. No significant associations were found for analyses of thalamus and nucleus accumbens volumes. CpG sites associated with hippocampus volume were significantly enriched within cancer-related genes and within regulatory elements containing the transcriptionally repressive histone H3K27 tri-methylation mark that is vital for stem cell fate specification. Cluster-based analyses revealed additional differentially methylated regions (DMRs) associated with hippocampal volume. DNA methylation at these loci affected expression of proximal genes involved in in learning and memory, stem cell maintenance and differentiation, fatty acid metabolism and type-2 diabetes. These DNA methylation marks, their interaction with genetic variants and their impact on gene expression offer new insights into the relationship between epigenetic variation and brain structure and may provide the basis for biomarker discovery in neurodegeneration and neuropsychiatric conditions.

## INTRODUCTION

Structural brain measures are important correlates of developmental and health outcomes across the lifetime. A large body of evidence has revealed age-related reductions in grey matter structures across the brain ^1^, notably in the hippocampus, which correlates with declining memory performance in older adults ^2, 3^. Recent findings from large-scale neuroimaging analyses within the ENIGMA consortium have revealed consistent patterns of cortical ^4, 5^ and subcortical ^5-8^ brain volume reductions across several neuropsychiatric disorders. Of all structures reported, the hippocampus was the most consistently and robustly altered, being smaller in major depressive disorder ^6^, schizophrenia ^7^, Attention deficit hyperactivity disorder (ADHD) ^8^ and Posttraumatic stress disorder (PTSD) ^9^. Other notable changes included volume reductions in the thalamus and NAcc in schizophrenia ^7, 8^.

Such differences in brain structure may fundamentally reflect the effects of genetic and environmental factors and their interplay, as suggested by the study of discordant monozygotic twins ^10^. DNA methylation is an epigenetic mechanism that may underlie gene-environment contributions to brain structure. It is under the influence of genetic ^11, 12^ and developmental ^12-14^ factors and plays an important role in brain development and disease, by regulating gene expression. DNA methylation is also a mechanism through which external stimuli, such as the environment, may contribute to expression of common diseases such as neurodegenerative disorders ^15^.

While efforts to identify genetic factors influencing brain structure have flourished in recent years ^16-18^, epigenetic studies of brain-related phenotypes remain very sparse. A considerable constraint is the need for a surrogate tissue for epigenetic studies of the living human brain. Crucially, initial reports have demonstrated that although DNA methylation patterns are largely tissue-specific, often differing between blood and brain ^19, 20^, there are also similarities ^21^ and blood DNA methylation shows promise as a biomarker for brain-related traits, including neuropsychiatric disorders ^22- 26^, cognitive ability ^27, 28^ and future psychopathology ^25^. However, only a few studies of small sample sizes have reported associations between blood DNA methylation and brain phenotypes ^25, 29, 30^.

Here, we built upon these findings and performed a large multisite epigenome-wide association study (EWAS) of structural brain volumes in 3,337 individuals from 11 cohorts. We focussed on analyses of the hippocampus, thalamus and NAcc – three disease-related subcortical regions of varying heritability ^17, 31^, and with large volumetric variability ^32^.

## MATERIAL AND METHODS

### Subjects and brain measures

The brain phenotypes examined in this study are from the ENIGMA analysis of high-resolution MRI brain scans of volumetric measures (full details in ^17^). Our analyses were focussed to mean (of left and right hemisphere) volumetric measures of three subcortical areas: the hippocampus, thalamus and nucleus accumbens, selected for their link to disease, different levels of heritability, and developmental trajectories. MRI brain scans and genome-wide DNA methylation data were available for 3,337 subjects from 11 cohorts (Supplementary Table 1).

### DNA methylation microarray processing and normalization

Blood DNA methylation was assessed for each study using the Illumina HumanMethylation450 (450k) microarray, which measures CpG methylation across >485,000 probes covering 99% of RefSeq gene promoters ^33^, following the manufacturer’s protocols.

Quality control procedures and quantile normalization were performed using the *minfi* Bioconductor package in R ^34^. Briefly, red and green channels intensities were mapped to the methylated and unmethylated status, and average intensities used to check for low quality samples. Initial quality assessment of methylation data was performed using the preprocessIllumina option. Principal component analyses (PCA) were performed using the singular value decomposition method, to identify methylation outliers based on the first four components. Samples with intensities more than 3 standard deviations away from the median were considered outliers and were removed. Intensities from the sex chromosomes were used to predict sex, and samples with predicted sex different from their recorded value were removed. Samples that were initially processed in batches were merged at this stage before further preprocessing. Stratified quantile normalization was then applied across samples The data were then normalized together using the *minfi* preprocessQuantile function ^35^. PCA of normalized beta values were used to control for unknown structure in the methylation data. Most cohorts estimated the cell counts for the 6 major cell types in blood (granulocytes, B cells, CD4+ T cells, CD8+ T cells, monocytes and NK cells) for each individual by implementing the estimateCellCounts function in *minfi*, which gives sample-specific estimates of cell proportions based on reference information on cell-specific methylation signatures. Other cohorts (i.e., NTR) measured cells counts directly.

### Epigenome-wide association analysis

Epigenome-wide association studies with volumes of the thalamus, hippocampus and NAcc were performed for each site separately. After normalization, probes on the sex chromosomes were filtered out (which are more difficult to accurately normalize), as were probes not detected (detection p-value > 0.01) in more than 20% of samples and probes containing a SNP (minor allele frequency ≥ 0.05) at the CpG or at the single nucleotide extension site.

We modelled association of DNA methylation and mean brain volumes in the hippocampus, thalamus and NAcc using linear regression analyses. Control variables included sex, age, age ^2^, intracranial volume, methylation composition (the first 4 principal components of the methylation data), and blood cell-type composition (first two components of estimated cell-type proportion) and depending on the sample and disease status (when applicable). For studies with data collected across several centres, dummy-coded covariates were also included in the model. Cohorts with family data (NTR, QTIM) performed association analyses using generalized estimating equation to control for familial relationship in addition to the other covariates. Our analyses focused on the full set of subjects, including patients, to maximise the power to detect effects. We also re-analysed the data excluding patients to ensure that the effects detected were not driven by disease.

The EWAS results from each site were uploaded to a central server for meta-analyses. Cross-reactive probes were further removed from the EWAS result files from each site, leaving 397,164 probes for subsequent analysis. Results from each cohort were meta-analysed by combining correlations across all 11 cohorts with fixed effect model, weighting for sample size ^36^. False discovery rates (FDR) were computed (correcting for the number of brain regions tested) and FDR < 0.05 was considered statistically significant.

### Epigenetic correlations

The results of the meta-analyses were further used to evaluate the similarities in epigenetic contributions, i.e. the epigenetic correlations, between the volumes of the three subcortical regions based on a procedure established for genetic correlations ^37^, with few adaptions. Due to the lack of a universal methylome reference, we used our largest cohort, IMAGEN, to calculate the methylation similarity score (MS-score, analogous to the LD-score) that reflects physical links between DNA methylation events (methylation clusters). MS-score was computed for each methylation probe, as the sum of its squared correlations with all probes within a given sliding window centered at the probe in question. To achieve stable estimations, we tested varied sizes of sliding window, e.g. 2, 3 and 5Mb, again comparably to what have been used for LD-score calculations. In addition, the standard deviation of epigenetic correlation was evaluated through bootstrapping process, which is supposed to provide a more reliable estimation of variance than jackknife under general conditions ^38^.

### Identification of differentially methylated regions (DMRs)

We identified DMRs by applying the *Comb-p* algorithm ^39^ on the meta-analysis of hippocampal volume. *Comb-p* adjusts *p-*values for genomic autocorrelation (ACF), identifies enriched regions of low *p-*values, and performs inference on putative DMRs using Sidăk multiple testing correction ^40^. The ACF distance was set to 500bp and the *p-*value threshold required for a DMR at *p* < 0.05. DMRs contained a minimum of 2 CpG sites.

### Functional and enrichment analyses

Gene annotation, gene-based test statistics and enrichment analysis were performed using the GREAT v3.0.0 ^41^ annotation tool with default parameters. For the annotations, CpGs were considered ‘within’ the regulatory region of a gene if they fell within a region including 5.0 kb upstream and 1.0 kb downstream of its transcription start site and extending in both directions to the nearest gene or up to 1000 kb max. All regions were based on human genome (hg19) coordinates. For gene-based tests and pathway analyses, GREAT was run against a whole genome background and results were considered significant if they exceeded the significance threshold for two measures of enrichment: one using a binomial test over genomic regions and one using a hypergeometric test over genes.

To test for enrichment for genomic regions found associated with hippocampal volume in our recent GWAS meta-analysis of hippocampal volume ^17^, we performed analyses based on MAGENTA ^42^, a computational tool designed for gene sets-based enrichment analyses with GWAS meta-analyses data as an input. To avoid “double dipping” in these analyses, we excluded the IMAGEN sample from the ENIGMA hippocampal volume meta-analysis, which we used as a dataset for known hippocampal volume SNPs (i.e., the ‘gene set’). We then tested for enrichment of this ‘gene set’ in the IMAGEN hippocampus EWAS results.

We modified the MAGENTA program to make it suitable for the analysis of DNA methylation data by first creating a ‘gene set’ of SNP regions by mapping SNPs to genomic locations, taking into account recombination hotspots. Adjacent regions with recombination rates lower than 10 were merged together. We then mapped CpG sites identified in the EWAS onto genomic regions if they fell within 100 kb of regions’ boundaries. Regions were scored based on p-values of the most significant CpG in the region. In addition, Šidák correction ^40^ was applied to correct for confounders such as gene size. Regions with significant enrichment were identified by permutation testing, using 5000 permutations. Two parameters were set to test for significant enrichment: *i*) the p-value threshold for selecting significant regions from the GWAS meta-analysis (GWAS thresholds of 5 x 10^-6^ and 5 x 10^-7^ were used) and *ii*) the cut-off threshold for each permutation: 90% and 99% cut-offs were used.

### Effects of methylation on gene expression

Effects of DNA methylation on gene expression were investigated in 631 subjects of the IMAGEN sample for which gene expression data were available. Total RNA was extracted from whole blood cells collected at the age of 14 using the PAXgene Blood RNA Kit (QIAGEN Inc., Valencia, CA). Following quality control of the total RNA extracted, labeled complementary RNA (cRNA) was generated using the Illumina^®^ TotalPrep™ RNA Amplification kit (Applied Biosystems/Ambion, Austin, TX). The size distribution of cRNA was determined through Bioanalyzer (Agilent Technologies, Santa Clara, CA) using the Eukaryotic mRNA Assay with smear analysis. Gene expression profiling was performed using Illumina HumanHT-12 v4 Expression BeadChips (Illumina Inc., San Diego, CA). Expression data were normalized using the mloess method ^43^. Expression data for genes mapping the top two CpG sites and DMRs associated with hippocampus volume. These included *BAIAP2* (probes ILMN_1705922, ILMN_1652865, ILMN_1699727, ILMN_2247226 and ILMN_2258749), *ECH1* (ILMN_1653115), *CMYA5* (ILMN_1805765) and its neighbouring genes *MTX3* (ILMN_1679071) and *PAPD4* (ILMN_1681845) genes, HHEX (ILMN_1762712) and *CPT1B* (ILMN_1791754). Expression data were log-transformed before analyses. For each DMR, a single DNA methylation factor was computed, taking into account methylation at all CpG sites within the DMR. Associations between gene expression and DNA methylation were measured using linear regressions with the first 4 principal components of the methylation data, sample batches, the first two components of estimated cell-type proportion, recruitment centres (dummy-coded) and sex as covariates.

### Methylation quantitative trait loci (mQTL)

To determine the relationship between genetic variation and CpG methylation levels, we searched for mQTLs in several datasets. First, we interrogated the ARIES dataset ^44^ that includes DNA methylation collected from peripheral blood (or cord blood) at five different time points across the life course from individuals in the Avon Longitudinal Study of Parents and Children (ALSPAC) ^45^. This dataset applied conservative multiple testing correction (p < 1 × 10^−14^) to identify between 24,262 and 31,729 sentinel associations at each time point.

We complemented this search using data from the combined Lothian Birth Cohorts (1921 and 1936), and the Brisbane Systems Genetics Study ^46^. The discovery and replication thresholds set in that study were*P* < 1 × 10^-11^ and *P* < 1 × 10^-6^, respectively, with both cohorts acting as a discovery (*P* < 1 × 10^-11^) and replication (*P* < 1 × 10^-6^) data set (only the most significant SNP for each CpG was considered).

### Expression quantitative trait loci (eQTL)

We used the Genotype-Tissue Expression (GTEx) database ^47^ to identify expression quantitative trait loci (*cis*-eQTLs; i.e., SNPs correlating with differential expression of neighbouring genes). This dataset, generated from ^48^ tissues from 620 donors, tests for significant SNPs-genes pairs for genes within 1Mb of input SNPs. The data described in this manuscript were obtained from the GTEx Portal (https://gtexportal.org/home/), Release: V7. It used FastQTL ^48^, to map SNPs to gene-level expression data and calculate q-values based on beta distribution-adjusted empirical p-values. A false discovery rate (FDR) threshold of <0.05 was applied to identify genes with a significant eQTL. The effect sizes (slopes of the linear regression) were computed in a normalized space (i.e., normalised effect size (NES)), where magnitude has no direct biological interpretation. They reflect the effects of the alternative alleles relative to the reference alleles, as reported in the GTEx database.

### Brain-blood methylation correlation

We interrogated a searchable DNA methylation database ^49^ (https://epigenetics.essex.ac.uk/bloodbrain/) generated from matched DNA samples isolated from whole blood and 4 brain regions (prefrontal cortex, entorhinal cortex, superior temporal gyrus, and cerebellum) from 122 individuals to establish the degree to which blood methylation levels at selected loci correlated with their brain methylation patterns. Correlations between blood and brain methylation levels at individual CpG loci were extracted and compared to Z-values from the hippocampal EWAS. An additional search was performed using data from blood and Brodmann areas 7, 10 and 20 from post-mortem samples of 16 individuals ^50^.

## RESULTS

### Associations of DNA methylation with subcortical volumes: analyses of individual CpG sites

We first investigated the association of DNA methylation at individual CpG sites in whole blood samples with the mean bilateral volumes of the hippocampus, thalamus and nucleus accumbens. Meta-analysis was applied by combining correlations across all 11 cohorts with fixed effect model, weighting for sample size. We identified 2 CpGs associating with volume of the hippocampus (Figure 1A; Supplementary Table 2) at an experiment-wide (correcting for the number of brain regions tested) false discovery rate (FDR) <0.05. The analyses of thalamus and NAcc volumes identified no CpG reaching the experiment-wide FDR threshold. Quantile-quantile (Q-Q) plots for the P-values of the analyses showed no evidence of P-value inflation. The CpGs associated with hippocampal volume explained each 0.9% of the phenotypic variance. Their effects were consistent across cohorts, with similar effect sizes for the cg26927218 site (*P*>0.1, Cochran’s *Q* test), while moderate heterogeneity in the magnitude, but not the direction of effects was noted for cg17858098 (Figure 1B). Effect sizes for analyses with and without patients across the 11 cohorts were very highly correlated (*r* ≥ 0.99) for CpGs with *P* < 1 x 10^-3^, indicating that these effects were unlikely driven by disease. These CpGs were annotated to the brain-specific angiogenesis inhibitor 1-associated protein 2 (*BAIAP2*) gene (also known as *IRSp53*; cg26927218) – encoding a synaptic protein whose expression in the hippocampus is required for learning, memory ^51^ and social competence ^52^– and to the enoyl-CoA hydratase-1 (*ECH1*; cg17858098), which encodes an enzyme involved in the β-oxidation of fatty acids ^53^.

**Figure 1:**
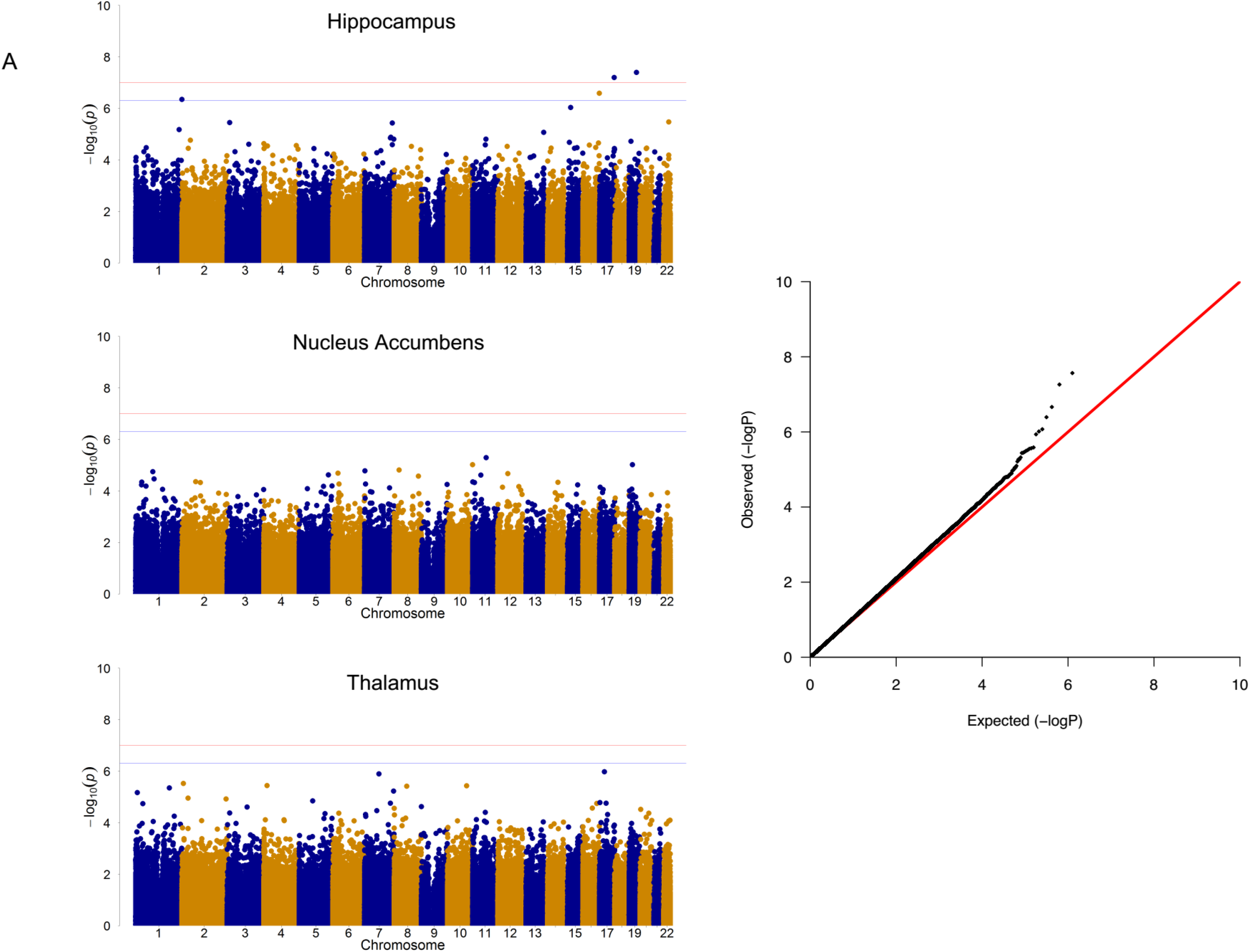
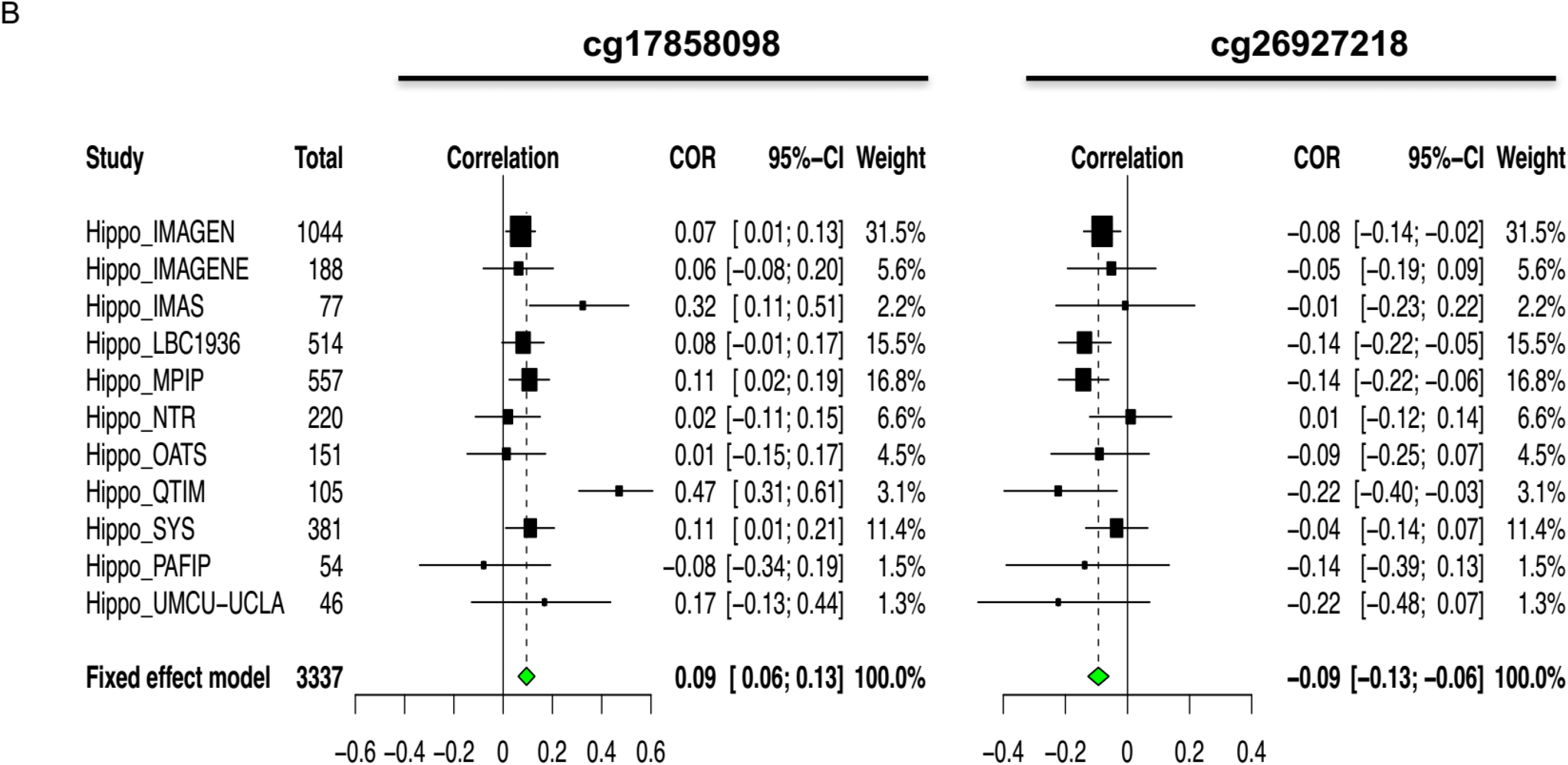
**A**, Manhattan plots *(left*) summarizing the association results for the hippocampus, thalamus and NAcc volumes. The red and blue lines represent the genome-wide FDR significance level (corrected for 3 brain regions) and non-corrected FDR significance level, respectively. Quantile-quantile plots (right) of multivariate GWAS of all traits (volumes of the hippocampus, thalamus and accumbens) show that the observed *P* values only deviate from the expected null distribution at the most significant values, indicating no undue inflation of the results. **B,** Forest plots show the effect (i.e., correlations between CpG methylation and hippocampus volume) at each of the contributing sites to the meta-analysis. The size of the dot is proportional to the sample size, the correlation level is shown on the *x* axis, and confidence interval is represented by the line.

### Functional annotation and enrichment analyses

To gain insight into functional relationships shared by genes mapping to differentially methylated CpGs associated with hippocampal volume, we performed enrichment analyses ^41^ on 340 CpGs associated with hippocampus volume at P < 5 x 10^-4^ (Supplementary Table 2). These CpGs showed effects specific for this structure rather than pleiotropic effects across the brain: most associations were unique to the hippocampus, a few shared with the thalamus and very few with the NAcc (Figure 2A). These closer epigenetic links between hippocampus and thalamus reflected closer correlations between their volumes (*r_H*T_* = 0.367, p = 5.78 x 10^-34^ and *r_H*N_* = 0.201, p = 8.36 x 10^-11^, for correlations of hippocampal volumes with thalamus and NAcc volumes, respectively).

**Figure 2:**
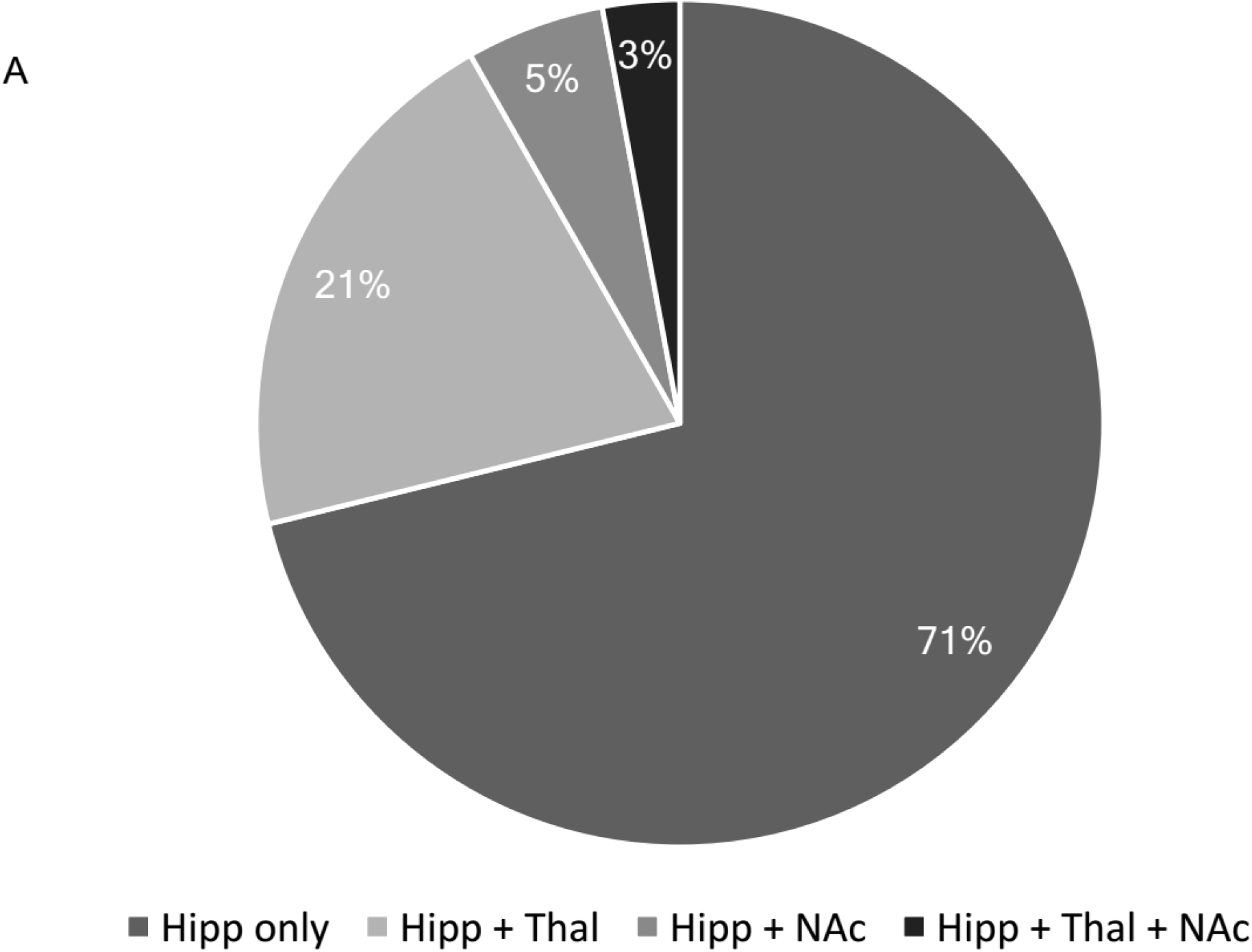
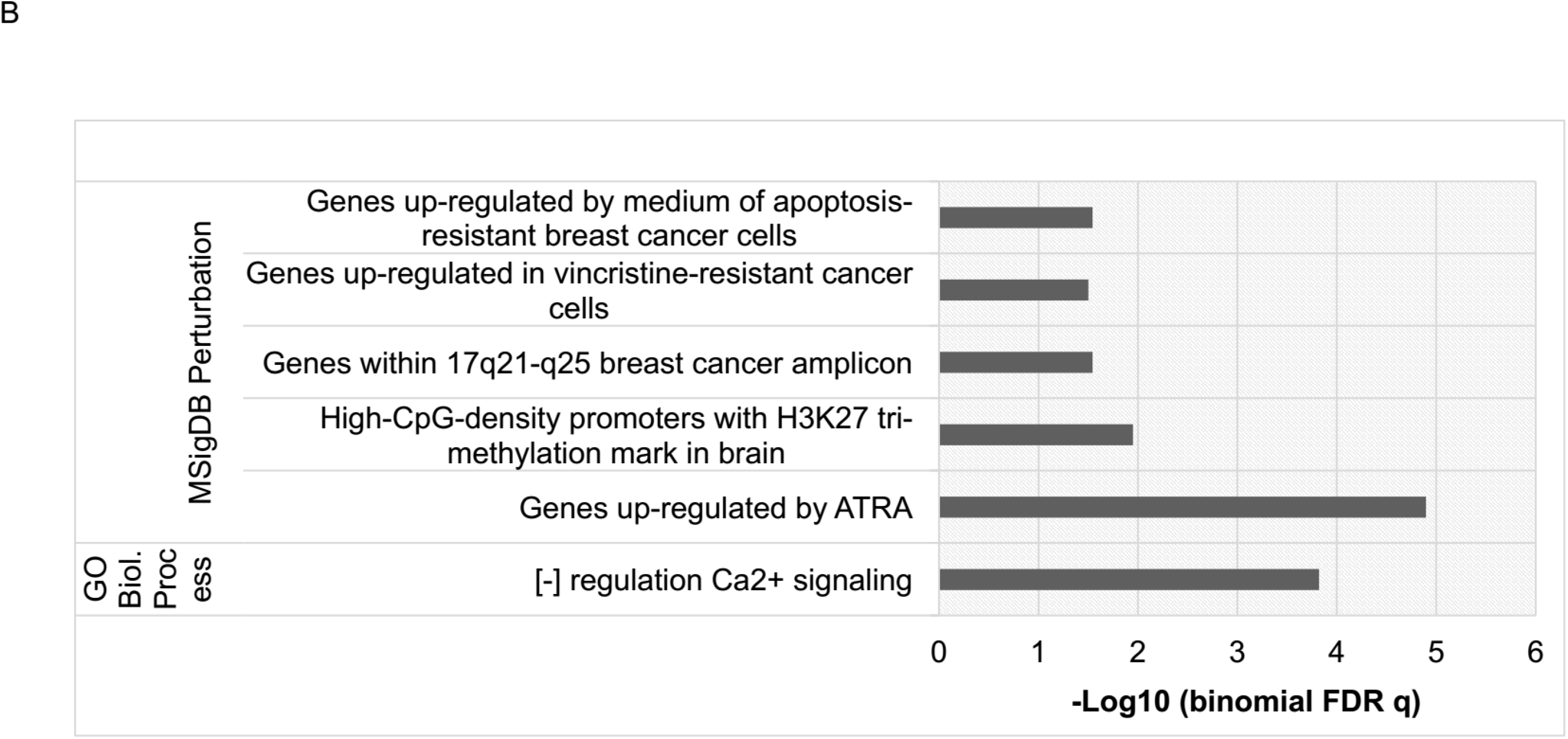
**A**, Pie chart of distribution of the 340 CpGs associated with hippocampus volume at P < 5x 10^-4^. The chart indicates the proportion of these CpG sites that are unique to the hippocampus or that are also associated (nominally, at p < 0.05) with the 2 other volumetric phenotypes investigated. In general, CpGs that influence other phenotypes than hippocampus volume have higher effect on thalamus than on NAcc volume. **B**, GREAT ontology enrichments of genes annotated to CpGs associated with hippocampus volume at P < 5x 10^-4^. Displayed are enriched terms from the Molecular Signatures Database (MSigDB) showing that genes annotated to CpGs associated with hippocampus volume are enriched for genes under epigenetic control and cancer-related genes.

Many genes annotated to these CpGs were related to cancer –including genes amplified or upregulated in cancers such as *BAIAP2* that lies within the 17q21-q25 breast cancer amplicon–, apoptosis and genes whose expression is affected by anti-cancer treatment (Figure 2B). In addition, there was significant overrepresentation of genes with high-CpG-density promoters carrying the histone H3K27 tri-methylation mark in the brain (n = 26). Such a feature is typical of key developmental genes ^54, 55^ targeted by the polycomb repressive complex 2 (PRC2) (Supplementary Table 3), a class of polycomb group proteins that by repressing gene expression via H3K27me3 is vital for maintenance of embryonic stem cell fate and cancer development. Accordingly, this set of genes was enriched for transcriptional regulators (n = 18 out of 26) including *TP73*, encoding a tumor suppressor protein that, by affecting neurogenesis, has a specific role in regulating hippocampal morphogenesis ^56^.

### Associations of DNA methylation with subcortical volumes: Cluster-based analyses

The analyses described above didn’t account for effects of DNA methylation clusters at regions formed by spatially correlated CpGs, which often occur within regulatory regions in the genome and are powerful means to control gene expression. Therefore, in the following analyses we studied the impacts of such DNA methylation clusters on the volumes of each subcortical region, i.e. the hippocampus, thalamus and NAcc. We first computed the methylation similarity score (MS-score) for each CpG within a given genomic region, that is analogous to the genetic LD-score, and that reflects the extent to which DNA methylation correlate between CpG sites in that region. We then regressed MS-scores with the squared Z-statistics derived the meta-analysis of each subcortical region and estimated the variance explained by this epigenetic information (analogous to the genetic variance ^37^) from the slope of the regressions. While we observed positive epigenetic variance for hippocampus (ρH^2^ = 0.060, p = 4.10 x 10^-3^, Supplementary Figure 1A) and thalamus (ρT^2^ = 0.044, p = 0.0303, Supplementary Figure 1B), a significant negative epigenetic variance was found for the NAcc (ρN^2^ = - 0.078, p = 2.42 x 10^-4^, Supplementary Figure 1C). This suggests that DNA methylation clusters contribute to the volumes of the hippocampus and –to a lesser extent– the thalamus. Conversely, mainly rare DNA methylation events or isolated CpG sites (with small MS-scores) may contribute to the volume of the NAcc ^37^. It is also notable that intercepts estimated in the above MS-score regressions were close to 1, ranging between 1.011~1.083 (Supplementary Table 4), suggesting generally small impact of confounding factors ^37^. Given the similarities in the epigenetic architecture noted above, we further characterised the epigenetic correlation between hippocampus and thalamus volumes. The estimated epigenetic covariance between these two traits was computed ^57^ (ρ_H*T_ = 0.0468; p = 2.27 x 10^-3^), and used to the estimate epigenetic correlations (r = ρ_H*T_/(ρ_H_*ρ_T_); Supplementary Table 5). These analyses revealed high epigenetic correlations (e.g., *r* = 0.91; s.d. = 0.235) between hippocampus and thalamus volumes, correlations significantly differing from 0 (t = 3.88, p = 4.37 x 10^-4^) but not from 1 (t = 0.38, p = 0.741), suggesting that hippocampus volume and thalamus volume share similar associations with methylation, consistent with the closer epigenetic links between hippocampus and thalamus reported in the section above. Of note, while the above results were generated with MS-scores based on a sliding window size of 3 Mb, very similar results were obtained with MS-scores calculated with window sizes of 2 Mb and 5 Mb (Supplementary Tables 3 & 4).

### Identification of Differentially Methylated Regions

Thus, we set out to identify such DNA methylation clusters (i.e., differentially methylated regions, DMRs) by applying the *comb-p* algorithm ^39^ to our epigenome-wide meta-analyses of hippocampal volume. Several DMRs significantly associated with the volume of hippocampus in the meta-analysed results (Šidák ^40^ corrected *P* < 0.05, number of consecutive probes ≥ 2; total numbers of DMRs = 20; Table 1). A DMR that included the cg26927218 site was identified, further supporting association of *BAIAP2* methylation with hippocampal volume. In addition to being identified from the meta-analysed data, some of these DMRs were identified in at least 2 cohorts, when analyses were run on EWAS results of each cohort separately, indicating that their association with brain volumes were unlikely to be due to chance. They were located within the cardiomyopathy associated gene 5 (*CMYA5*; this DMR is subsequently referred to as DMR1), encoding an expression biomarker for diseases affecting striated muscle ^58-61^ and possibly a schizophrenia risk gene ^62^; the hematopoietically expressed homeobox (*HHEX;* DMR2) gene, encoding a homeobox transcription factor controlling stem cells pluripotency and differentiation in several tissues ^63-67^, and a well-known risk loci for type 2 diabetes ^68^, as well as the carnitine palmitoyltransferase 1B (*CPT1B;* DMR3) gene, encoding a rate-limiting enzyme in the mitochondrial beta-oxidation of long-chain fatty acids, whose expression enhances reprogramming of somatic cells to induced pluripotent stem cells ^69^, cancer cell self-renewal and chemoresistance ^70^. There was a significant degree of correlation of DNA methylation at these DMRs, being higher between DMR1 and DMR3, than DMR1 and DMR2 (r = 0.155, p = 7.30 x 10^-8^ and r = 0.147, p = 2.91 x 10^-7^, respectively). These DMRs were also taken forward for further analyses.

**Table 1:**
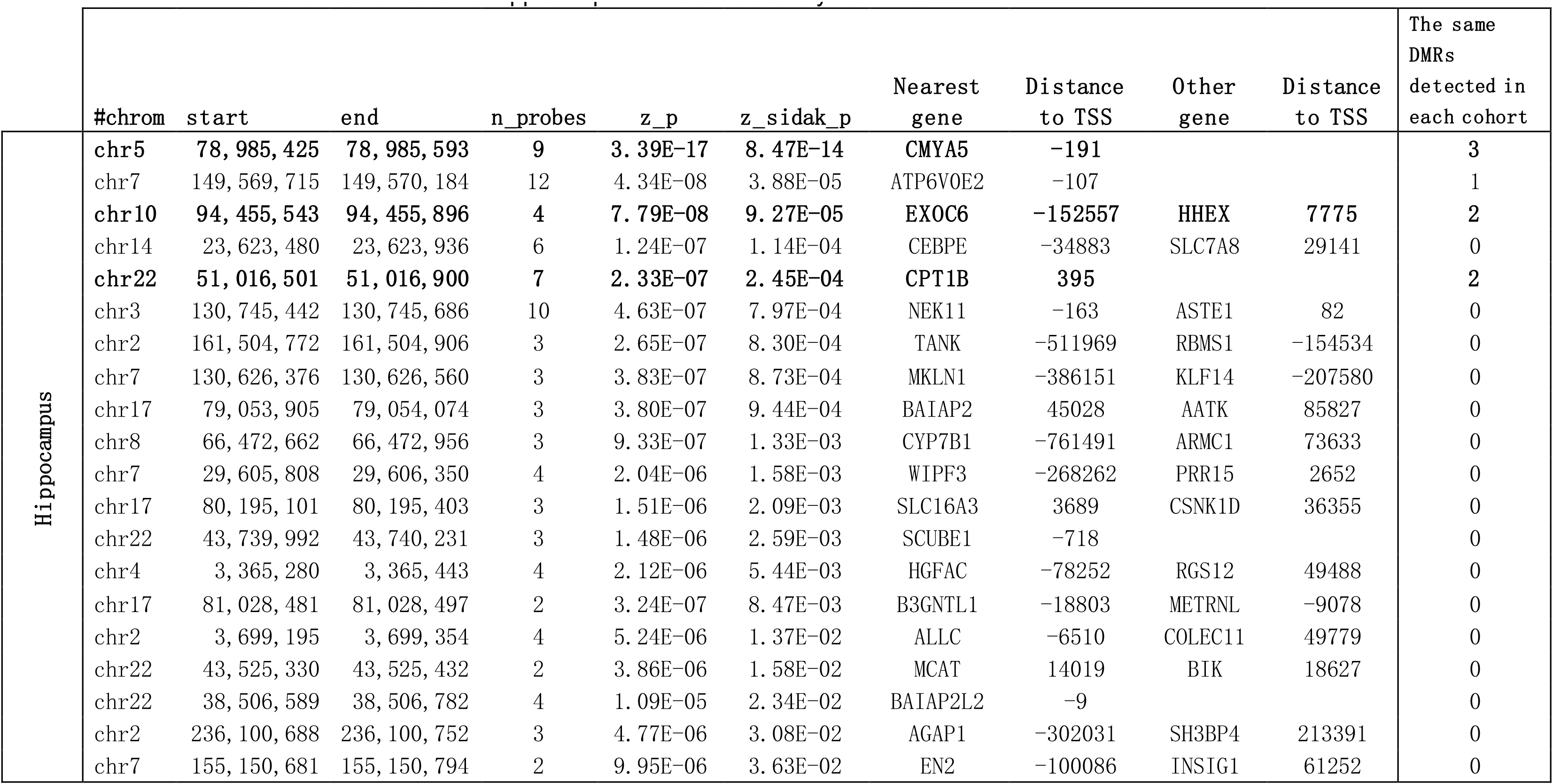
List of DMRs identified from the the hippocampus EWAS meta-analysis results

### Effects of differential methylation on gene expression

We measured the impact of DNA methylation on expression of neighbouring genes (*cis*-effects) in the IMAGEN sample. Methylation at most loci affected gene expression, with the effects of DMRs being larger than that of individual CpGs (i.e., cg26927218). Several isorforms are expressed from *BAIAP2* and isoform-specific effects were observed for cg26927218; methylation at this locus correlated with increased expression of the short isoform for *BAIAP2* (β = 0.016, p = 5 x 10^-3^; Figure 3A). There were no significant effects of cg17858098 on *ECH1* mRNA levels (β = −0.008, p = 0.201). Given the correlations between DMRs noted above, we controlled for methylation at the other 2 DMRs when testing for effects of a given DMR on gene expression. As shown in Figure 3B, DMR1 methylation had no effect on expression of *CMYA5* (β = −0.227, p = 0.492), tending instead to have contrasting effects on expression of neighbouring genes (β = −0.410, p = 0.039 and β = 0.554, p = 0.019 for *PAPD4* and *MTX3*, respectively). Methylation at DMR2 increased expression of its closest gene, *HHEX* (β = 0.351, p = 0.020). Methylation at DMR3 had strong effects on expression of the adjacent *CPT1B* gene (β = 1.670, p = 2.55 x 10^-59^). *Trans-*effects were also noted for this DMR, as it associated with increased expression of *PAPD4* (β = 0.724, p = 1.21 x 10^-7^), a gene adjacent to DMR1.

**Figure 3:**
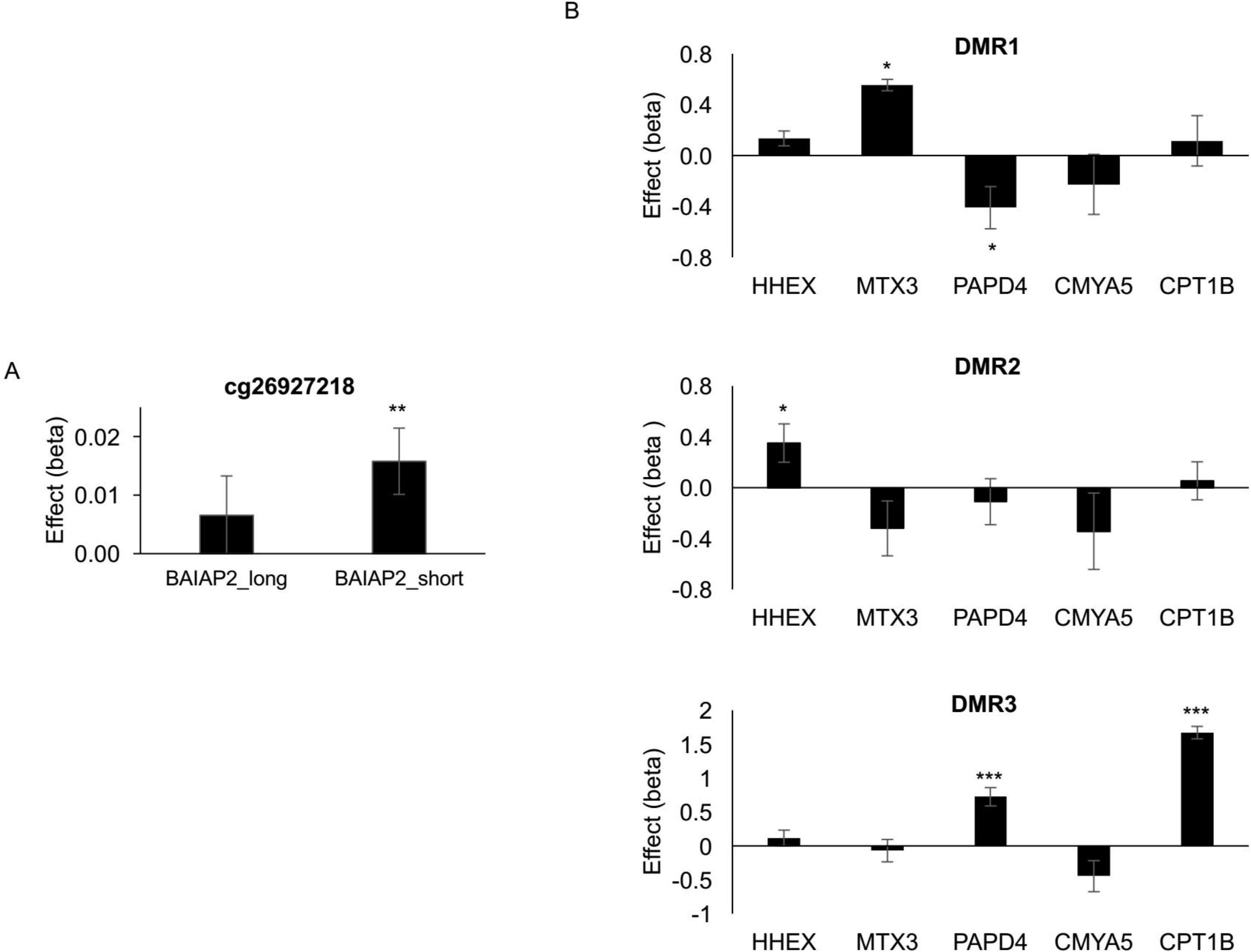
Analyses of top CpG (**A**) and DMRs (**B**) demonstrate effects of DNA methylation on gene expression in 631 subjects from the IMAGEN sample. In the DMR analyses, linear regression analyses tested relationship between methylation at the listed DMR and expression of *HHEX*, *MTX3, PAPD4, CMYA5* and *CPT1B*, controlling for methylation at the other 2 DMRs. Results represent unstandardized coefficients ± S.E.M. *, p < 0.05; **, p < 0.01; ***, p < 0.001.

### Correlations of DNA methylation between blood and brain

To investigate if the above findings would remain relevant for the brain, we tested if interindividual variation in whole blood predicted interindividual variation in the brain at the differentially methylated loci by conducting two blood–brain comparisons. First, we compared methylation levels at these sites in blood and brain tissues (blood, prefrontal cortex, entorhinal cortex, superior temporal gyrus and cerebellum) sampled from the same individuals (N = 75) using the blood–brain DNA methylation comparison tool ^49^ (Supplementary Table 6). There was no significant correlation between blood and brain methylation levels at the individual CpGs sites (cg26927218 –*BAIAP2*– and cg17858098 –*ECH1*). On the other hand, inter-individual variation in whole blood was a moderate predictor of inter-individual variation in the brain for DMR1 and DMR3 (strongest correlations: *r* = 0.54, p = 1.20 x 10^-6^ and *r* = 0.59, p = 2.37 x 10^-8^, respectively). For DMR2, correlations were more varied with the strongest correlation in the superior temporal gyrus (*r* = 0.37, p = 9.68 x 10^-4^). Correlations were stronger in cortical brain regions than in the cerebellum. The degree of this co-variation predicted the associations of DNA methylation at DMRs with hippocampal volume (Supplementary Table 6, Supplementary Figure 2), suggesting that mirroring blood DNA methylation within these DMRs in brain tissues may be what drives association of their association with hippocampal volume.

Another comparison between methylation in blood and in other brain regions – Brodmann area (BA)7 (parietal cortex); BA10 (anterior prefrontal cortex) and BA20 (ventral temporal cortex)– using BECon ^50^ revealed similar patterns (Supplementary Figure 3). For DMR1, there were moderate correlations between blood and BA7 methylation at all CpGs (r = 0.13-0.47) and between blood and BA10 for most CpGs (r = 0.13-0.30). For DMR3, correlations between blood and brain methylation were strong in all areas (r = 0.37-0.86), while the degree of correlations varied at DMR2 ranging from −0.35 to 0.34, depending on the CpG site and the brain area.

### Genetic contributions to differential DNA methylation associated with hippocampal volume

Given that genetic factors may underlie the correlations between DNA methylation in different tissues, we searched for methylation QTLs in two datasets. A search in the ARIES mQTL database ^45^ identified several SNPs associated with methylation at the DMR1 and DMR3 loci (Supplementary Table 7). The strongest mQTLs, rs131758 and rs4441859 affected methylation such that the A-allele at these SNPs associated with increased methylation at DMR3 and DMR1, respectively. These effects were replicated in two other datasets (Supplementary Table 7). Remarkably, eQTL analyses indicated that these alleles correlated with expressions of *CMYA5* and *CPT1B*, albeit differently. While the effects of the rs4441859_A allele were tissue-specific, the rs131758_A allele increased *CPT1B* expression in all tissues, including the brain (Supplementary Table 7 & Supplementary Figure 4).

Furthermore, we considered whether there was a significant overlap between DNA methylation differences identified in this study and SNPs associated with hippocampal volume. To test this, we used the recent genome-wide association studies of hippocampal volume conducted by ENIGMA ^17^ (excluding the IMAGEN data; GWAS association thresholds P < 5 x 10^-6^ and P < 5 x 10^-7^) as a dataset for significant hippocampal SNP regions, adapting MAGENTA ^42^, the gene sets-based enrichment analysis tool for GWAS data to the analysis of methylation data. SNPs were merged into genomic regions that were then examined for overlap with DNA methylation identified in hippocampal EWAS performed in the IMAGEN sample. These analyses revealed significant overlap between DNA methylation loci and SNP loci influencing hippocampal volume (Supplementary Table 8).

## DISCUSSION

In this large epigenome-wide meta-analysis we identified for the first time differentially methylated CpG sites and genomic regions whose levels of DNA methylation are predictive of variation in hippocampal volume. We further demonstrate the potential of blood to discover epigenetic biomarkers for the living human brain. Methylation at these sites affect the expression of genes required for hippocampal function, stem cell fate and function and genes involved in metabolic regulation. The observation that, at the identified DMRs, DNA methylation variation in blood mirrors that of brain tissues helps us generate hypotheses as to how modifiable factors such as diet and lifestyle may contribute to some of the impairments associated with diabetes and neurodegenerative conditions^71^.

Changes in hippocampal volumes are hallmarks of brain development predictive of cognitive deficits generally associated with aging and neurodegeneration. While large hippocampal volume is linked with good memory and cognitive function, hippocampal atrophy is associated with the development of a range of neurodegenerative ^72^ and neuropsychiatric disorders ^6^, ^7^, ^8 9^. Modifiable factors such as obesity, exercise, stress and medication can reduce or increase the size of the hippocampus throughout life ^72^. Collectively, our findings support these observations, pointing to associations of hippocampal volume with epigenetic mechanisms controlling cell fate and fatty acid metabolism, as discussed below:

First, two of the top hits identified (*CPT1B* and *ECH1*) encode key enzymes involved in β–oxidation of fatty acids. These enzymes act on the same pathway, CPT1B being necessary for the transport of long-chain fatty acids into the mitochondria and ECH1 for a key step in their β–oxidation. Fatty acids (notably the omega-3 polyunsaturated fatty acids) benefit brain development and healthy brain aging by modulating neurogenesis and protecting from oxidative stress throughout the lifespan ^73^. More specifically, like cancer stem cells ^70^, neural precursors in the hippocampus and subventricular zone require fatty acid oxidation for proliferation ^74^. This led to the proposition that abnormalities in brain lipid metabolism contribute to hippocampal dysfunction in AD by their ability to suppress neurogenesis at early stages of disease pathogenesis ^75^. Accordingly, fatty acid metabolism in the brain seems to be closely related to the pathogenesis of Alzheimer’s disease ^76^. Given such functional coupling between lipid metabolism, proliferation of progenitor cells in the brain and other tissues, including cancer ^74^ and neurodegeneration, findings that *ECH1* expression may serve as marker for AD ^77^ are not surprising. Likewise, CPT1B-dependent fatty acid β-oxidation has been found to be critical for breast cancer stem cell self-renewal and chemoresistance ^70^..

Further links between metabolism and hippocampal volume were suggested by our identification of a region annotated to a replicated risk locus for T2D (*HHEX*) ^68^. The metabolic alterations observed in T2D may induce cognitive dysfunction ^78^ by exacerbating declines in hippocampal volumes associated with aging ^79^ and AD pathology ^80^, a process to which *HHEX* may contribute ^81^. This is supported by findings that genetic variations within the *HHEX* gene region may underlie the association of T2D with AD, with the *HHEX* rs1544210_AA genotype interacting with diabetes to increase the risk of dementia and AD by more than four-fold ^81^. Furthermore, individuals with diabetes who carry the *HHEX* rs1544210_AA genotype tend to have significantly smaller hippocampal volumes than those without these conditions ^81^. This genotype is significantly associated with decreased *HHEX* expression in several tissues, as determined by analysis of the Genotype-Tissue Expression database (data not shown) supporting and complementing our findings that variations which increase expression of this gene (such as *HHEX* DNA methylation or the rs1544210_GG genotype) associate with larger hippocampal volume.

Finally, our analyses revealed sets of cancer-related genes and genes carrying the repressive tri-methylation of histone H3 at lysine 27 (H3K27me3 mark) that silences key developmental genes. This epigenetic mark typically targets genes controlling stem cell renewal, which are commonly deregulated in cancer ^82-84^. Intriguingly, the HHEX transcription factor was recently recognized as a regulator of stem cell fate ^63-67^ and an oncogene that enabled H3K27me3-mediated epigenetic repression of tumour suppressor genes ^63^. This role of *HHEX* in transcriptional control links epigenetic mechanisms controlling stem cell fate to pathological processes underlying metabolic and neurodegenerative diseases.

DNA methylation at most loci had clear, albeit distinct effects on gene expression. Notable transcript-specific effects were observed for cg26927218 on *BAIAP2*. The cg26927218 locus is located in a DNase I hypersensitive site, characteristic of regions actively involved in transcriptional regulation ^85^, within a consensus DNA binding sequence for the MYC associated factor X (MAX) – a transcription factor controlling cell proliferation, differentiation, and apoptosis. MAX belongs to a class of transcription factors that recognize CpG-containing DNA binding sequences, only in their unmethylated form ^86, 87^. Thus, methylation at cg26927218 may affect expression of the *BAIAP2* short variant by directly interfering with the function of this transcription factor. A role for the region surrounding cg26927218 in transcriptional regulation is further supported by findings showing that a genetic variant (rs8070741) near cg26927218 enhances cortical expression of the *BAIAP2* short variant ^88^.

Besides the hippocampus, none of the other two subcortical structures investigated generated significant results. This may reflect a unique role of the hippocampus in brain development, possibly related to it being a site of neurogenesis. These findings are also consistent with the relative heritability of the different subcortical structures, indicating higher twin-based heritability estimates for larger (hippocampus and thalamus) compared to smaller (NAcc) subcortical structures but overall low SNP-based heritability ^17^. This supports the model according to which a substantial fraction of the heritability of complex traits is due to epigenetic variation ^89^. Our analyses on genetic contributions to DMRs’ effects also suggest that epigenetic control is partially modulated by genetic variations, which is further suggested by the overlap between GWAS and EWAS of hippocampal volume.

In conclusion, we have identified DNA methylation at several loci that affect hippocampus volume, which indicate for the first time possible mechanistic pathways by which modifiable and metabolic factors might contribute to the pathology of neurodegenerative diseases. A clear limitation is the small number of cohorts for which both MRI and DNA methylation data are available, we nonetheless provide a rigorous roadmap that should encourage larger and more extensive future studies. Our work demonstrates the usefulness of combining peripheral epigenetic markers and neuroimaging measures to discover epigenetic factors that may predict brain status and illustrate the unrivalled opportunity to understand the biological mechanisms through which modifiable factors contribute to common human diseases.

## Acknowledgements

**ENIGMA:** The study was supported in part by grant U54 EB020403 from the NIH Big Data to Knowledge (BD2K) Initiative, a cross-NIH partnership, and by NIH grant R56 AG058854 to the ENIGMA World Aging Center.

**IMAGEN**: This work received support from the following sources: the European Union-funded FP6 Integrated Project IMAGEN (Reinforcement-related behaviour in normal brain function and psychopathology) (LSHM-CT-2007-037286), the Horizon 2020 funded ERC Advanced Grant ‘STRATIFY’ (Brain network based stratification of reinforcement-related disorders) (695313), ERANID (Understanding the Interplay between Cultural, Biological and Subjective Factors in Drug Use Pathways) (PR-ST-0416-10004), BRIDGET (JPND: BRain Imaging, cognition Dementia and next generation GEnomics) (MR/N027558/1), the FP7 projects MATRICS (603016), the Innovative Medicine Initiative Project EU-AIMS (115300-2), the Medical Research Council Grant ‘c-VEDA’ (Consortium on Vulnerability to Externalizing Disorders and Addictions) (MR/N000390/1), the National Institute for Health Research (NIHR) Biomedical Research Centre at South London and Maudsley NHS Foundation Trust and King’s College London, the Bundesministeriumfür Bildung und Forschung (BMBF grants 01GS08152; 01EV0711; eMED SysAlc01ZX1311A; Forschungsnetz AERIAL 01EE1406A, 01EE1406B), the Deutsche Forschungsgemeinschaft (DFG grants, SM 80/7-2, SFB 940/2, NE 1383/14-1), the Medical Research Foundation and Medical research council (grant MR/R00465X/1) and by NIH Consortium grant U54 EB020403, supported by a cross-NIH alliance that funds Big Data to Knowledge Centres of Excellence. Further support was provided by grants from: ANR (project AF12-NEUR0008-01 – WM2NA, and ANR-12-SAMA-0004), the Fondation de France, the Fondation pour la Recherche Médicale, the Mission Interministérielle de Lutte-contre-les-Drogues-et-les-Conduites-Addictives (MILDECA), the Fondation pour la Recherche Médicale (DPA20140629802), the Fondation de l’Avenir, Paris Sud University IDEX 2012; the National Institutes of Health, Science Foundation Ireland (16/ERCD/3797), U.S.A. (Axon, Testosterone and Mental Health during Adolescence; RO1 MH085772-01A1), the Swedish Research Council (Vetenskapsrådet), the Swedish Research Council for Health, Working Life and Welfare (FORTE), the Swedish Research Council FORMAS (grant number 259-2012-23), the 111 Project (NO.B18015), the NSFC (81801773), the key project of Shanghai Science & Technology (No.16JC1420402), the Shanghai Municipal Science and Technology Major Project (No.2018SHZDZX01), ZHANGJIANG LAB, and the Shanghai Pujiang Project (18PJ1400900).

**LBC1936**: We thank the cohort participants and team members who contributed to these studies. Phenotype collection in the Lothian Birth Cohort 1936 was supported by Age UK (The Disconnected Mind project). Methylation typing was supported by the Centre for Cognitive Ageing and Cognitive Epidemiology (Pilot Fund award), Age UK, The Wellcome Trust Institutional Strategic Support Fund, The University of Edinburgh, and The University of Queensland. Analysis of the brain images was funded by the Medical Research Council Grants G1001401, 8200, and MR/M01311/1. The imaging was performed at the Brain Research Imaging Centre, The University of Edinburgh (http://www.bric.ed.ac.uk), a centre in the SINAPSE Collaboration (http://www.sinapse.ac.uk). The work was undertaken by The University of Edinburgh Centre for Cognitive Ageing and Cognitive Epidemiology (http://www.ccace.ed.ac.uk), Part of the cross council Lifelong Health and Wellbeing Initiative (Ref. G0700704/84698); Funding from the Biotechnology and Biological Sciences Research Council and Medical Research Council (MR/K026992/1) are gratefully acknowledged. The Scottish Funding Council contributed support through the SINAPSE Collaboration.

**MPIP**: The MPIP sample comprises patients included in Munich Antidepressant Response Signature study and the Recurrent Unipolar Depression (RUD) Case-Control study. We acknowledge Rosa Schirmer, Elke Schreiter, Reinhold Borschke and Ines Eidner for image acquisition and data preparation. We thank Stella Iurato for supporting quality control procedures of the methylation measurements. The MARS project was supported by the German Federal Ministry of Education and Research (BMBF) through the NGFN and NGFN-Plus programs (FKZ 01GS0481), the Molecular Diagnostics program (FKZ 01ES0811), the Research Network for Mental Diseases program (FKZ 01EE1401D) and by the Bavarian Ministry of Commerce. This work was also funded by the German Federal Ministry of Education and Research (BMBF) through the Integrated Network IntegraMent (Integrated Understanding of Causes and Mechanisms in Mental Disorders), under the auspices of the e:Med Programme (grant # 01ZX1314J to EB).

**NTR**: The NTR study was supported by the Netherlands Organization for Scientific Research (NWO), MW904-61-193 (Eco de Geus & Dorret Boomsma), MaGW-nr: 400-07-080 (Dennis van ‘t Ent), MagW 480-04-004 (Dorret Boomsma), NWO/SPI 56-464-14192 (Dorret Boomsma), the European Research Council, ERC-230374 (Dorret Boomsma), and Amsterdam Neuroscience.

**OATS:** We thank the OATS participants and their supporters for their time and generosity in contributing to this research. We acknowledge the contribution of the OATS Research Team (https://cheba.unsw.edu.au/project/older-australian-twins-study). OATS was supported by an Australian National Health and Medical Research Council (NHMRC)/Australian Research Council Strategic Award (ID401162) and a NHMRC Project Grant (ID1045325). This research was facilitated through Twins Research Australia, a national resource in part supported by a NHMRC Centre for Research Excellence Grant (ID: 1079102).

**PAFIP:** This work was supported by the Instituto de Salud Carlos III (PI14/00639 and PI14/00918), MINECO (SAF2010-20840-C02-02 and SAF2013-46292-R) and Fundación Instituto de Investigación Marqués de Valdecilla (NCT0235832 and NCT02534363). No pharmaceutical company has financially supported the study.

**QTIM**: Acknowledgements: We thank the twins and singleton siblings who gave generously of their time to participate in the QTIM study. We also thank the many research assistants, radiographers, and IT support staff for data acquisition and DNA sample preparation.

Funding Acknowledgements: National Institute of Child Health & Human Development (RO1 HD050735); National Institute of Biomedical Imaging and Bioengineering (Award 1U54EB020403-01, Subaward 56929223); National Health and Medical Research Council (Project Grants 496682, 1009064 and Medical Bioinformatics Genomics Proteomics Program 389891).

**SYS**: This work was funded by the Canadian Institutes of Health Research, Canadian Foundation for Innovation and Heart and Stroke Foundation of Canada.

**UMCU**: The UMCU cohort was supported by the Netherlands Organization for Health Research and Development (ZonMw) numbers 908-02-123 and 917-46-370 (Hilleke Hulshoff Pol), and High Potential program (Hilleke Hulshoff Pol) of the Utrecht University.

## Author contributions

<u>Manuscript Writing and Editing</u>: S.D. wrote the manuscript; B.R., E.B.B., F.H., G.S., J.L., N.F., P.M.T., P.S., T.C-R., T.J., T.J.E., T.P., U.B., V. C., V.F., and Z.P. edited the first draft; all authors critically reviewed the manuscript

<u>Cohort Principal Investigators</u>: A.S., B.C-F., D.A., D.I.B., E.B.B., G.S., H.B., I.J.D., J.T., L.G.A., M.J.W., M.W., P.R.S., P.S.S., R.A.O., V.C. and Z.P.

<u>Imaging data acquisition</u>: A.d.B, D.v.E, E.A., F.N., G.I.Z., H.F., J-L.M, J.J., J.L., J.T., K.L.M., L.T.S., M.E.B., M.H., P.M.T., P.S., R.B., T.B., T.P., V.F., W.C. and W.W.

<u>Epigenetic data acquisition</u>: A.F.M., A.T., D.S., G.B., J.L., J.S., J.v.D., J.T., K.A.M., N.G.M., P.S.,T.C-R., T.J., Y.L. and Z.P.

<u>Data analysis</u>: A.O., B.R., C.C., J.L., J.S., J.v.D, K.S., M.L., N.A., N.J., R.J., R.R-S.,S.D., T.C-R. and T.J.

---

**Supplementary Figure 1:** MS-score plots for hippocampus (A), thalamus (B) and NAcc (C) volumes. The vertical axis represents mean scores in MS-score quantiles and the horizontal axis the mean χ^2^ statistic of variants in each quantile. Colors correspond to regression weights, with red indicating large weight. The red line is the MS score regression line.

**Supplementary Figure 2:** Relationship between blood vs. brain correlation and association with hippocampal volume. The x-axis represents the effect (z-score) of individual CpGs within the listed DMR on hippocampal volume. The y-axis shows the corresponding correlation between DNA methylation in blood versus brain in 4 brain areas 49 at these CpGs. Generally, stronger effects are observed for CpGs sites whose methylation levels are highly correlated in at least one tissue.

**Supplementary Figure 3:** Comparison between DNA methylation in blood and in three brain regions (BA7, BA10 and BA20) in paired samples from 16 individuals ^50^. Metrics shown for CpG sites composing each of the 3 DMRs, include spearman correlation values of methylation between blood and the listed brain region, methylation variability in blood and brain samples and average methylation change with cell composition adjustment.

**Supplementary Figure 4:** Expression quantitative trait loci analyses showing effects of rs4441859 and rs131758 genotypes on *CMYA5* and *CPT1B* expression in tissues from 620 donors from the Genotype-Tissue Expression (GTEx) database ^47^. Effects fulfilling the FDR threshold of ≤0.05 are highlighted in red.

